# The trichotomy of pneumococcal infection outcomes

**DOI:** 10.1101/370007

**Authors:** Alexis Erich S. Almocera, Gustavo Hernandez-Mejia, César Parra-Rojas, Esteban A. Hernandez-Vargas

## Abstract

The successful elimination of bacteria such as *Streptococcus pneumoniae* from a host involves the coordination between different parts of the immune system. Previous studies have explored the effects of the initial pneumococcal load (bacterial dose) on different representations of innate immunity, finding that pathogenic outcomes can vary with the size of the bacterial dose. However, others yield support to the notion of dose-independent factors contributing to bacterial clearance. In this paper, we seek to provide a deeper understanding of the immune responses associated to the pneumococcus. To this end, we formulate a model that realizes an abstraction of the innate-regulatory immune host response. Stability and bifurcation analyses of the model reveal the following trichotomy of pneumococcal outcomes determined by the bifurcation parameters: (i) *dose-independent clearance*; (ii) *dose-independent persistence*; and (iii) *dose-limited clearance*. Bistability, where the bacteria-free equilibrium co-stabilizes with the most substantial steady-state bacterial load is the specific result behind dose-limited clearance. The trichotomy of pneumococcal outcomes here described integrates all previously observed bacterial fates into a unified framework.

## 1. Introduction

The pneumococcus (*Streptococcus pneumoniae*) is a bacterial pathogen associated with pneumonia, otitis media (ear infections), and life-threatening conditions such as sepsis or bacteremia (blood poisoning) and bacterial meningitis (brain infection) [1]. The pneumococcus is notably a coinfective pathogen with the influenza virus, contributing to enhanced morbidity and mortality [2, 3, 4, 5]. According to the World Health Organization (WHO), the pneumococcus is the causative agent behind 16% of mortalities in children under five years of age, and the deaths of 920,136 children in 2015. Diseases caused by the pneumococcus are mostly common among children and the elderly, as well as individuals with a compromised immune system [6, 7, 8]. Current immunization programs may have desirable impacts [9, 10]; however, they remain challenged by antibiotic-resistant serotypes [8, 9, 11, 12]. These challenges emphasize the need for more research and development towards the control and eradication of *S. pneumoniae*.

Briefly speaking, the infection of the pneumococcus begins with entry of pneumococcal particles through the nasal cavity, followed by adherence to epithelial cells (colonization), and concluding with invasive disease [1]. An extensive collection of studies identified a diverse range of bacterial and environmental factors that contribute to the infection, including enzymes, binding regions, and capsule structures— see, *e.g.*, [13] and [14] for a review. However, here we are interested in the kinetics of *S. pneumoniae* and interactions of the pathogen with first responders, comprising innate immune responses and regulatory mechanisms. To this end, mathematical models provide conceptual frameworks for studying immune responses and changes in bacterial load.

Previous mathematical models investigate the effects of initial bacterial load (dosage or inoculum size) on the successful bacterial clearance by the innate responses. Model formulations assume the phagocytes to form a single group [15], separate phagocytes according to whether they actively engulf pneumococcus [16], or differentiate phagocytes into neutrophils and macrophages [17, 18]. However, these models did not consider the cascading effect of immune responses. Smith et al. [19] formulated three models fitted to bacterial load in mice to investigate the coordination of innate immune cells. This coordination is described by the following cascade of innate responses: alveolar macrophages, neutrophils, and monocyte-derived macrophages. Each model predicts that the bacteria will be cleared in small doses and sustain persistent levels in high doses; we refer to this phenomenon as *dose-limited clearance*. Besides the modeling and experimental work in [19], separate studies identified drug-specific effects [20], genetic variations in either the host or the pathogen [21, 22], and biological switches (for Toll-like receptors and bacteremia threshold) [23] as factors in the pneumococcal outcome in addition to inoculation size. In light of the models reviewed here, we may conceptualize a general form of the innate-regulatory response.

The phenomenon of dose-limited clearance is comparable to the behavior of a bistable system. For a broad range of nonlinear dynamical systems, a simple case of bistability follows from the coexistence of two asymptotically stable equilibrium points. Hence, a typical solution approaches one equilibrium point or the other, depending on the initial state [24]. Malka et al. [25] constructed a one-dimensional equation describing bacterial kinetics and clearance by constant densities of neutrophils. In a specific region of the parameter space, the model exhibits bistability under moderate neutrophil densities. That is, three steady states appear, where the bacteria-free value is mutually stable with the maximum. The intermediate value is unstable and serves as a clearance threshold comparable to Smith et al. [19]. This observation indicates the inadequacy of neutrophils to clear more massive bacterial loads. Furthermore, hysteresis accompanies bistability: when neutrophils decrease to critically small levels and then return to the original state, the clearance of sufficiently small bacterial population suddenly changes to an irreversible persistence event. Notably, the bistability phenomenon agrees with published data from a series of bactericidal experiments [26], suggesting that bistability is a plausible mechanism for fulminant infection. In this paper, we formulate a model of three ordinary differential equations, which generalizes the innate-regulatory immune response to *S. pneumoniae*. The generalized immune response unifies associated mechanisms into abstract components.

We organize the paper as follows. In Section 2, we introduce our model and estimate parameters by fitting the model to murine experimental data in [27]. The experiments in this study revealed that some but not all mice reached undetectable pneumococcal levels after 16 hours post-infection (hpi). Our analysis in Section 3 reveals a bistability event comparable to [25]. Moreover, our model predicts three possible outcomes: clearance, persistence, and dose-limited clearance in the bistability case. The first two outcomes are independent of the bacterial dose size. One threshold parameter controls the window of bistability, while another dictates the predicted outcome. Section 4 concludes this paper with a discussion.

## 2. Mathematical model

We establish a mathematical model to represent the global panorama of the bacterial (*B*) interaction with the host and the corresponding immune response which in this case is characterized by the innate (*M*) and regulatory (*N*) immune responses. The model reads as follows:

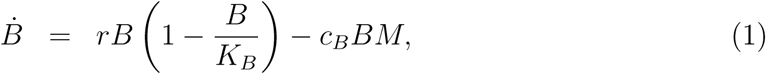

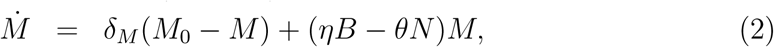

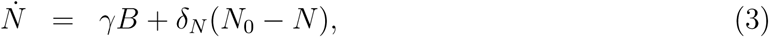

where the dot denotes the derivative with respect to a time variable *t*, *i.e.*,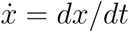, *x* = *B, M, N*. Here, the bacteria (*B*) proliferates logistically at a maximum rate *r* with a tissue carrying capacity of *K*_*B*_, given in colony forming units per milliliter (CFU/mL). The clearance of free bacteria occurs at a rate *c*_*B*_ per cell and is assumed to result from the innate immune response. *M* is assumed to evolve with a constant rate *η* due to bacterial presence. We consider a constant decline rate of *M* given by *θ*, which is influenced by the immune regulatory activity. The elimination rate of *M* is given by *δ*_*M*_. Furthermore, the regulatory process is favored by bacteria at a rate *γ* and its inhibition rate is given by *δ*_*N*_. The initial immune response levels are *M*_0_ (innate) and *N*_0_ (regulatory). We assume a constant replenishment of the innate (*δ*_*M*_ *M*_0_) and regulatory (*δ*_*N*_ *N*_0_) responses. Figure 1 is a conceptual diagram of the model (1)–(3).

**Figure 1:**
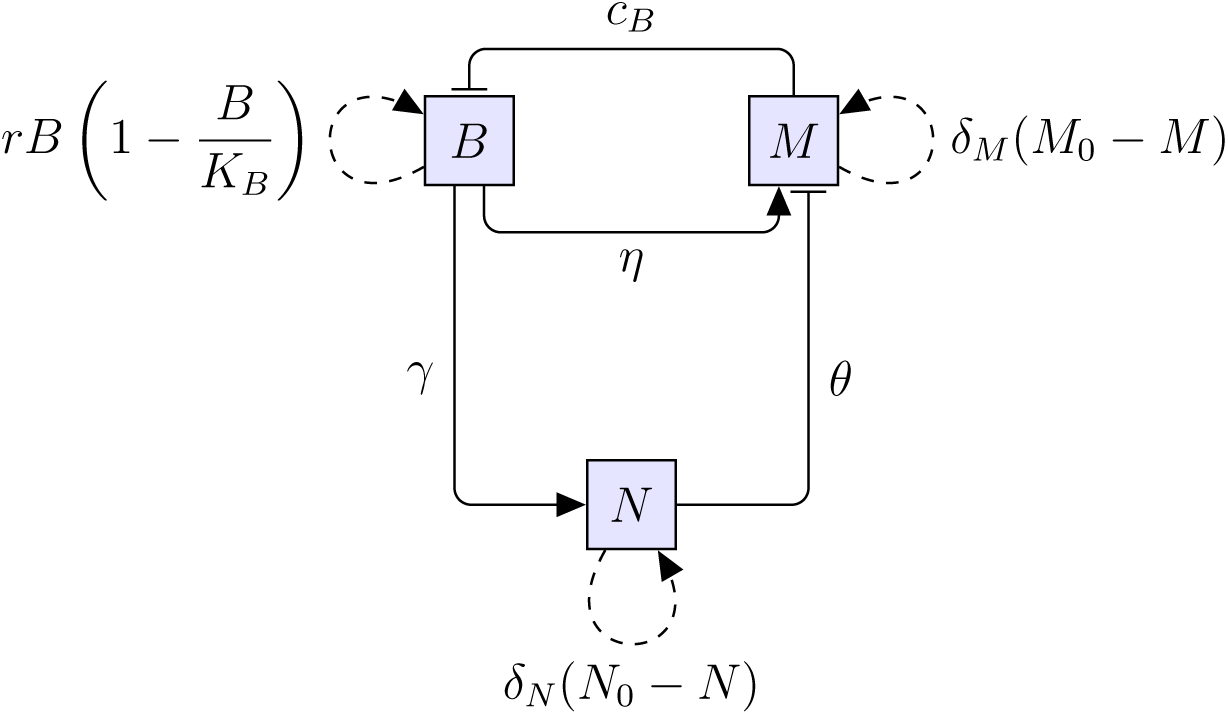
Schematic diagram of the model (1)–(3). The state variables for bacterial load (*B*), innate (*M*) and regulatory (*N*) response levels are depicted in shaded squares. Solid lines with arrowheads indicate bacterial activation of innate and regulatory responses, associated with constant rates *η* and *γ*, respectively. Solid lines ending in bars denote the following inhibitory effects: regulatory levels inhibit innate response growth at a constant rate *θ*, and the innate response controls bacterial growth at a constant clearance rate *c*_*B*_. Dashed loops indicate the replenishment of a given state variable according to its corresponding growth term.

### Experimental data

In the experiment of Duvigneau *et al.* [27], C57BL/6J wildtype mice were intranasally infected with a sub-lethal dose of 1*×*10^6^ CFU of the *S. pneumoniae* strain TIGR4 (T4). After the infection, the bacterial load in the bronchoalveolar lavage was measured at different time points, namely at 1.5, 6, 18, 26 and 31 hpi. Figure 2 depicts the experimental data of bacterial load in the lungs of the infected mice. Note that the experiment presents two outcomes, bacterial clearance (a) and persistence (b). We explore the interaction of bacteria with the host immune response through the combination of data and the proposed model (1)–(3) for the two scenarios. For each set of data in Figure 2, we separately estimated the majority of parameters. The remaining ones were set as described below.

**Figure 2:**
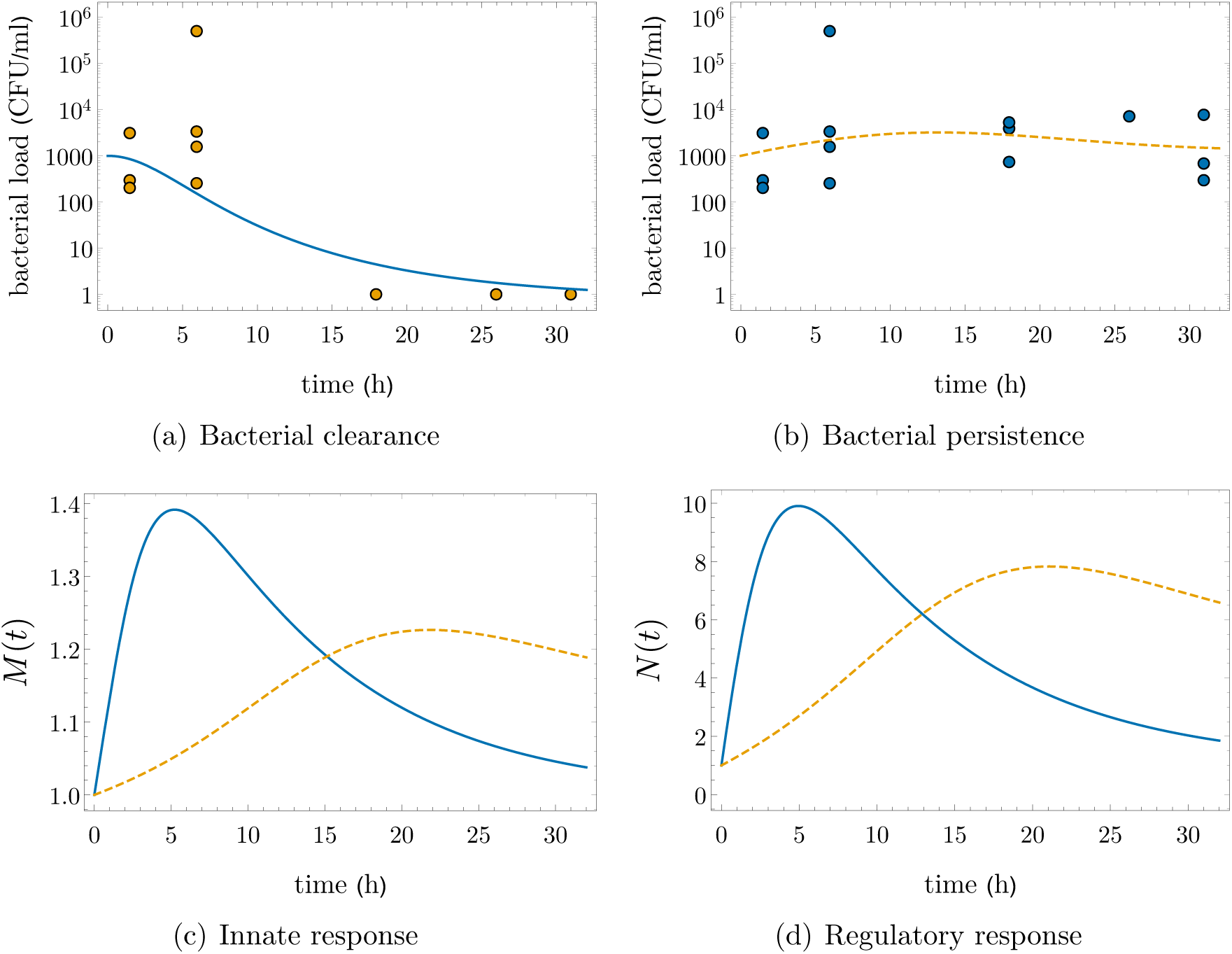
Dynamics of bacterial load and immune response. **Top:** bacterial data (circles) and fitted dynamics for (a) the clearance (blue, solid line) and (b) the persistence (orange, dashed line) scenarios. Note that in both panels the data at 1.5 and 6 hpi is the same. **Bottom:** (c) Innate and (d) regulatory immune responses for both bacterial behaviors: clearance (blue, solid line) and persistence (orange, dashed line).

### Parameter estimation

According to Smith et al. [19] and Duvigneau et al. [27], the bacterial growth rate *r* = 1.13 h^-1^ and the carrying capacity *K*_*B*_ = 2.3*×*10^8^ CFU/mL stand for single *S. pneumoniae* infection. Also, following the criteria from Smith et al. [19], the *S. pneumoniae* inoculum size (*B*_0_) for simulations is set to 1000 CFU/mL. Several elimination rates of immune actors such as alveolar macrophages, cytokines as interleukin-1 and tumor necrosis factor-*α*, neutrophils, and monocyte-derived macrophages are reported in [19] for the specific dynamical models there developed. In contrast, our model considers all innate response components as a single global response rather than as separate actors. The same philosophy is considered for the regulatory response. Under these considerations, we set the innate and regulatory inhibition rates as 0.1, a general-response value. In addition, for the global regulatory response, we set *M*_0_ = 1 due to the constant supply of innate agents, such as alveolar macrophages [19, 27]. A constant regulatory action is also considered assuming *N*_0_ = 1. We would like to remark that different assumptions of initial values and elimination rates would only rescale the fitted parameters, but not affect the mechanistic insights from the model selection procedures.

To fit the remaining parameters, *i.e.*, *c*_*B*_, *η*, *θ* and *γ*, we minimized the mean squared difference (MSD) between the model output and the experimental bacterial measurement, both on logarithmic scale. The model equations were solved in Python using the numerical integration routines of the SciPy library [28]. The minimization of the MSD was also performed with SciPy using the Differential Evolution algorithm [29]. Separately, we fitted each of the datasets to uncover the parameter values that yield either the persistence of the bacterium or its elimination. The fitted values are shown in Table 1, while Figure 2 shows the resulting dynamics of the model (1)–(3) for each case. Innate and regulatory responses are plotted in fold-change, Figure 2(c)–(d). The dynamics of the clearance case shows a marked bacterial elimination after 16 hpi—see Figure 2(a). Note that for the clearance scenario, both immune responses present a large marginal increase during the first hours post infection; this effect is more pronounced in the regulatory response, reaching a 10-fold increase before 7 hpi—see Figure 2(d). In contrast, the persistent bacteria appears to provoke a sluggish action of the immune response, making the regulatory and innate responses peak after 20 hpi, where the regulatory effect is almost reaching an 8-fold increase—see panels (c) and (d) of Figure 2.

**Table 1:** Fitted parameter values for the different bacterial infection outcomes. The sets of four parameters were independently fitted based on bacterial data from either the persistence or clearance scenarios.

| Parameter | Bacteria clearance | Bacterial persistence |
| --- | --- | --- |
| $c_B$ | 1.13096 | 0.97000 |
| $\gamma$ | 0.00376 | $2.85546 \times 10^{-4}$ |
| $\eta$ | $1.3461 \times 10^{-4}$ | $8.07453 \times 10^{-6}$ |
| $\theta$ | $1.3982 \times 10^{-13}$ | $3.50118 \times 10^{-13}$ |

## 3 Model analysis

We nondimensionalize the model (1)–(3) to simplify computations. Let us introduce the following dimensionless variables

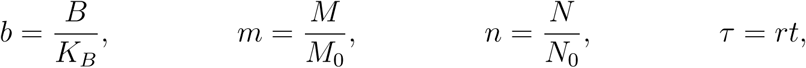

where *τ* is a rescaled time variable. This transformation scales the bacterial load with the carrying capacity, and the immune response levels with their respective initial values. Define the following dimensionless parameters:

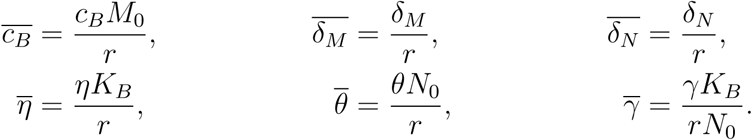

Then the model (1)–(3) has the dynamically equivalent dimensionless form

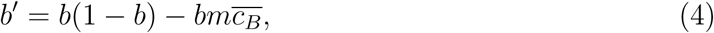

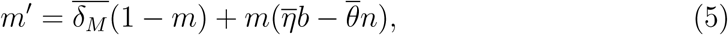

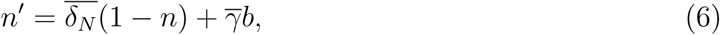

where the prime denotes the derivate with respect to the rescaled time *τ*.

To determine the local stability of system (4)–(6), we denote a point in statespace by its coordinates (*b, m, n*). Then (4)–(6) admits the unique equilibrium point

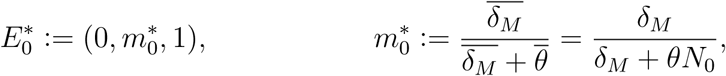

corresponding to steady state values 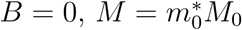 and *N* = *N*_0_. Let

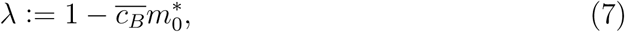

and denote the Jacobian matrix of (4)–(6) by *J* (*b, m, n*). Then the following result is evident from the eigenvalues of 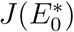, which take values 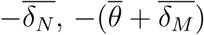 and *λ*.

### Theorem 1.

*The dimensionless system* (4)*–*(6) *admits the unique equilibrium point 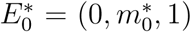 where b* = 0*. Moreover, 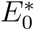 is asymptotically stable if λ <* 0 *and is a saddle point when λ >* 0.

We consider *λ* as our bifurcation parameter, and determine equilibrium points of the form *E*^*\**^ = (*b*^*\**^, *m*^*\**^, *n*^*\**^) where *b*^*\**^ *>* 0. To this end, we introduce the map

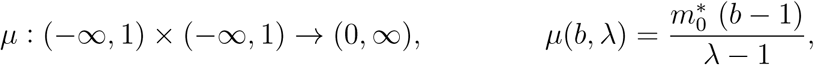

and let

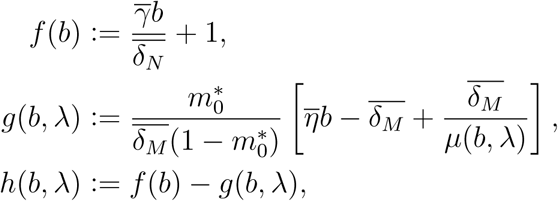

for *b <* 1 and *λ <* 1. We are interested in the roots of *h*(·, *λ*) in the open interval (0, 1) which are later determined to be the values of *b*^*\**^. To this end, we first establish the following monotone and concave properties, where 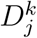 denotes the *k*th partial derivative with respect to the *j*th variable and 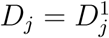.

### Lemma 1.

*The following properties hold:*

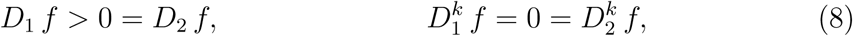

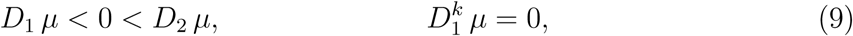

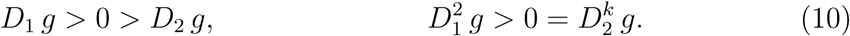

*where k≥*2*. Thus, f is a strictly increasing linear function, and g*(*b, λ*) *is strictly increasing and concave upwards in b while being strictly decreasing in λ. Moreover, 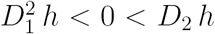 and 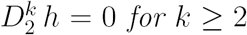 for k ≥* 2, *i.e., h is concave downwards in b and is a strictly increasing linear function in λ.*

*Proof.* The properties in (8) follow directly the definition of *f*, from which *f* is strictly increasing and linear. To establish (9), we have

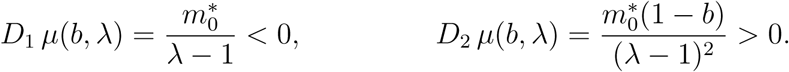

Consequently, *D*_1_ *µ*(*b, λ*) is independent of *b*, and 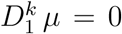 for *k ≥* 2. Thus, we compute *D*_1_ *g* and 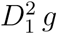 as follows:

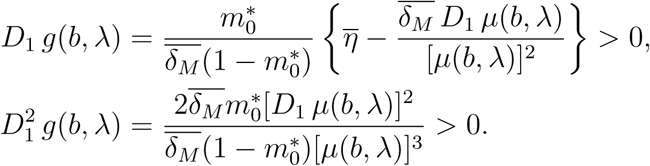

Finally, we have *D*_2_ *h* = *-D*_2_ *g* due to *D*_2_ *f* = 0, and we obtain

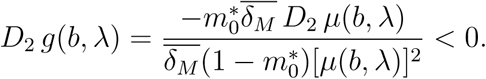

Since (*D*_2_ *µ*)*/µ*^2^ expands to a function independent of *λ*, we have 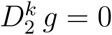 for *k≥*2. Therefore, the properties in (10) are true; in particular, *g* is strictly increasing and concave upwards in *b*, while being strictly decreasing in *λ*. The results for *h* = *f-g* follow from (8) and (10).

The results of *h* in Lemma 1 yields the following properties. First, the root of *D*_1_ *h*(·, *λ*) is uniquely given by a certain 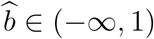 from which

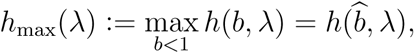

that is, 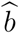 is the unique maximizer of *h*(·, *λ*). Furthermore, we have

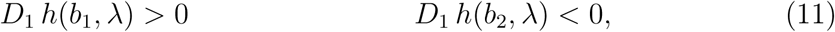

for arbitrary values 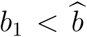 and 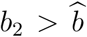. The function *h*_max_ is linear with positive slope because 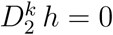 for *k ≥* 2. Hence, we define the unique root of *h*_max_ as

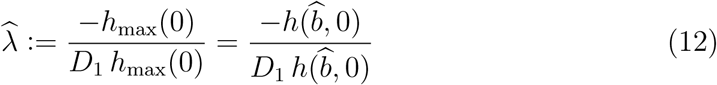

from Maclaurin expansion. That is, 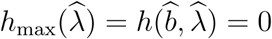.

Now, let

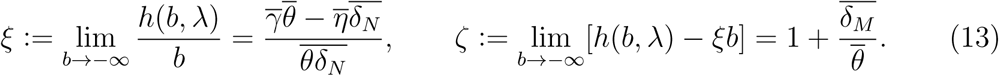

Then the curve *y* = *h*(*b, λ*) is asymptotic to the line *y* = *ξb* + *ζ* as *b→ -∞.* Moreover, the coefficients *ξ* and *ζ* of the asymptote line allow us to write

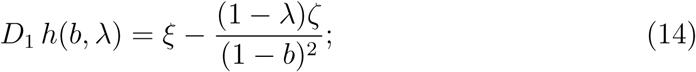

from this, we derive

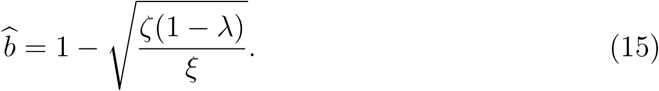

In the following lemma, we determine some limiting behavior on the function *h* and its first derivative.

### Lemma 2.

*For* 0 *< b <* 1, *we have h*(*b, λ*) *< bD*_1_ *h*(0, 0)*. Moreover,*

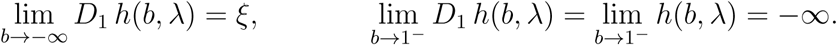

*Proof.* Recall that *h* strictly increases in *λ* (Lemma 1), and note that the curve *y* = *h*(*b,* 0) is tangent to the line *y* = *h*(0, 0) + *b D*_1_ *h*(0, 0) = *b D*_1_ *h*(0, 0) at *b* = 0. Hence by the concavity of *h*(·, *λ*), we have

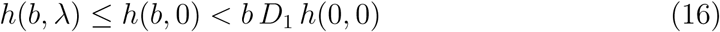

for *b >* 0. Passing the limit to (16) as *b →* 1^*-*^, we have lim_*b→*1_*-h*(*b, λ*) = *-∞*. We complete the proof by passing limits to (14) where *b → -∞* and *b →* 1^*-*^.

We establish that the values of *b*^*\**^ are roots of *h*(·, *λ*) in the interval (0, 1) on which *f >* 0. From now on, we denote the smallest and largest positive roots by 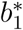 and 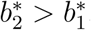, respectively.

### Lemma 3.

*The function h*(·, *λ*) *has at most two distinct roots, namely 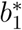 and 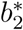. If 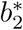 exists, then 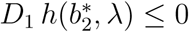. If 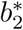 also exists, then 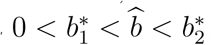, from which*

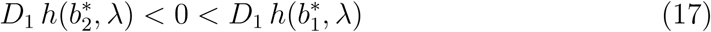

*and λ <* 0*. Moreover, the following equations are equivalent:*

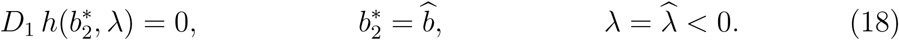

*Proof.* Rolle’s theorem asserts that a real-valued differentiable function with two distinct roots attains a local maximum or minimum at a point between the roots. Thus, a continuously differentiable function with at least three roots has no fixed concavity. Since *h*(·, *λ*) is concave downwards by Lemma 1, no more than two roots exist for *h*(·, *λ*), which are 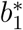 and 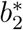.

Recalling from Lemma 2 that *h*(*b, λ*) *→ -∞* as *b →* 1^*-*^, suppose that 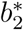 exists so that *h*(·, *λ*) has no root in the interval 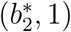. Assuming 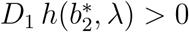 implies that *h*(*b, λ*) is initially positive then approaches *-∞*, as *b* increases from 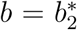 to *b* = 1. However, a contradiction arises with 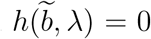 for some 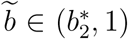. Thus, 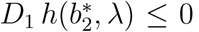. If 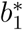 additionally exists, then 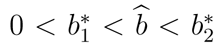 by Rolle’s theorem and the uniqueness of 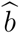 as the root of *D*_1_ *h*(·, *λ*). We obtain (17) by taking 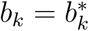 for each *k* in (11). Furthermore,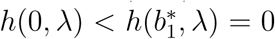. Since *h*(0, *λ*) has the same sign with *λ*, we have *λ <* 0.

Assuming that 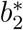 exists, 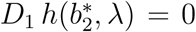 if and only if 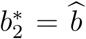 because 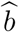 is the unique root of *D*_1_ *h*(·, *λ*). In such case, *h*(·, *λ*) strictly increases over the interval 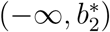, hence 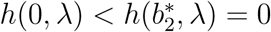 and *λ <* 0. Now, 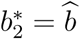 implies

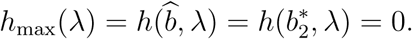

Conversely, *h*_max_(*λ*) = 0 implies that either 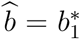 or 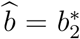. We must have 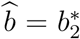 because the existence of 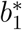 necessitates 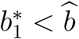 as shown above. Finally, *h*_max_(λ) = 0 if and only if 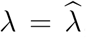 by the uniqueness of 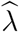 as the root of *h*_max_. Therefore, the equations in (18) are equivalent.

We now establish the existence of roots for the function *h*(·, *λ*) in (0, 1). In particular, we show that the existence of both 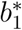 and 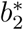 depends on the value of *D*_1_ *h*(0, 0), which from (14) is given by

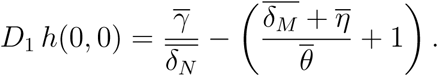

We may alternatively write

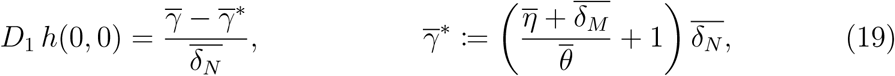

to frame our results with 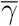, which is associated with the proliferation response of interferon growth due to bacterial stimuli. The following result establishes the existence of roots for the function *h*(·, *λ*). This result is illustrated in Figure 3.

**Figure 3:**
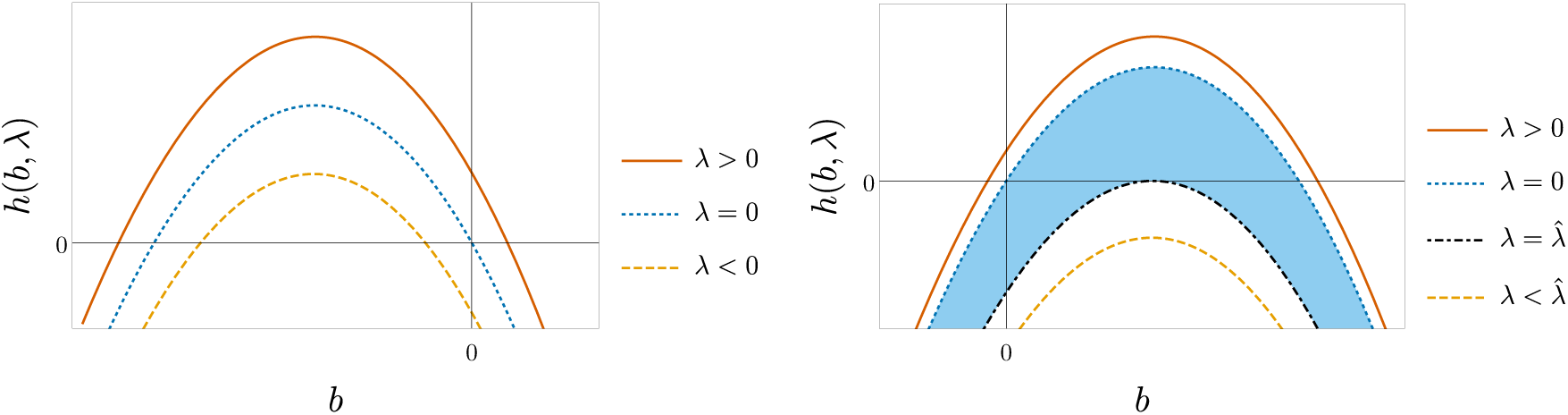
The function *h*(·, *λ*) depicted at different values of *λ*. In the left panel, 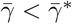 for which a unique positive root exists if and only if *λ >* 0. The right panel considers the case where 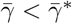. The shaded region identifies the family of curves generated by *h* where 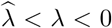, for which two distinct roots exist. The region is bounded by the bifurcation values *λ* = 0 and 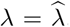. No root exists when 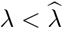 (bottom curve) and exactly one positive root exists for *λ >* 0 (top curve).

### Theorem 2.

*If λ >* 0, *then 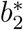 is the unique root of h*(·, *λ*) *in* (0, 1)*. If λ ≤*0, *then h*(·, *λ*) *admits roots in* (0, 1) *only if D*_1_ *h*(0, 0) *>* 0*. In this case, 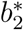 is the unique root whenever λ* = 0*. Moreover, the following trichotomy holds:*

(i) *Both 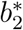 and 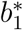 exist for 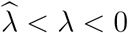*

(ii) *Only 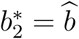 exists for 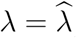 and*

(iii) *Neither 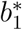 nor 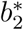 exists when 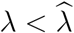.*

*Proof.* Recall that *h*(0, *λ*) is equal in sign to *λ*. If *λ >* 0, then 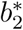 is a root of *h*(·, *λ*) in (0, 1) by the intermediate value theorem, because *h*(*b, λ*)*→ -∞* as *b →* 1^*-*^ from Lemma 2. Appealing to Lemma 3 where *λ >* 0, we arrive at the uniqueness of 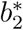 as the root in (0, 1).

Now, assume that *λ ≤* 0. If *D*_1_ *h*(0, 0) *≤* 0, then it follows from Lemma 2 that

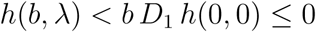

for *b >* 0. Thus, it is necessary that *D*_1_ *h*(0, 0) *>* 0 for *h*(·, *λ*) to have a root in the interval (0, 1). In this case, we infer from similar arguments as Lemma 3 that 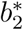 is the unique root whenever *λ* = 0. Observe that *D*_1_ *h*(0, *λ*)*→ -∞* as *λ→ -∞*, so that we may choose an integer *n >|λ|* such that *D*_1_ *h*(0, *-n*) *<* 0. The maximizer of *h*(*·,-n*) is negative by virtue of (11) where *λ* =*-n* and *b*_2_ = 0, as does the global maximum due to

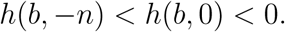

for *b <* 0. By contrast, *h*(*b,* 0) has a positive maximizer and a positive global maximum. Considering the continuity of the maximizer and global maximum of *h*(·, *λ*) as functions of *λ*, it follows that 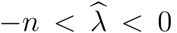. Now, recall that 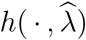 is maximized at 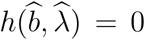. The desired trichotomy holds by comparing *h*(*b, λ*) with 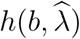 and *h*(*b,* 0), and appealing to the linear increasing property of *h* in *λ* (Lemma 1); this can be associated with the shaded region bounded by 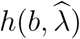 and *h*(*b,* 0), located in the right panel of Figure 3.

### Corollary.

*Given our bifurcation parameter λ, the only positive equilibrium points for the dimensionless model* (4)*–*(6) *are of the form*

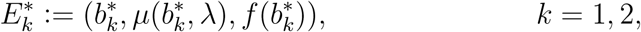

*Where 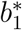 and 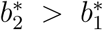 are the smallest and the largest positive roots of h*(·, *λ*), *respectively. If λ >* 0, *then only 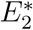 exists. If λ ≤* 0 *then positive equilibrium points exist only if D*_1_ *h*(0, 0) *>* 0 *and 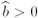. In this case, the following trichotomy holds: both 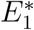 and 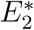 exist if 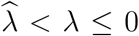 only 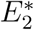 exists where 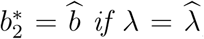 and no positive equilibrium point exists when 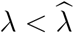.*

*Proof.* Consider a positive equilibrium point 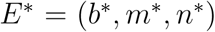. Then the following equations hold:

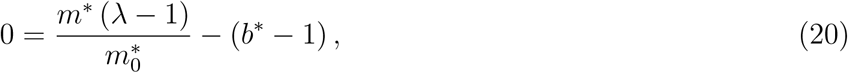

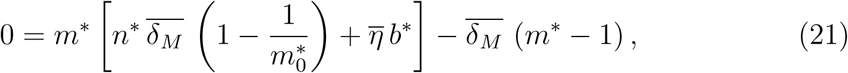

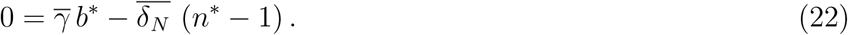

We rewrite equations (20) and (22) into *m*^*\**^ = *µ*(*b*^*\**^, *λ*) and *n*^*\**^ = *f* (*b*^*\**^), respectively. Thus, solving (21) in *n*^*\**^ after evaluating *m*^*\**^ yields *n*^*\**^ = *g*(*b*^*\**^, *λ*). Consequently, *h*(*b*^*\**^, *λ*) = *f* (*b*^*\**^)*-g*(*b*^*\**^, *λ*) = 0 and *b*^*\**^ is either 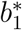 or 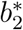 by Lemma 3. Thus, *E*^*\**^ is either 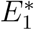 and 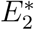. For each *k*, the equilibrium point 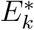 exists if and only if 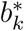 exists. Therefore, the existence 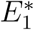 and 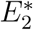 follows from Theorem 2.

Now, let

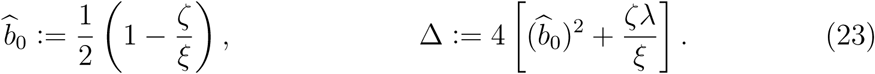

Then we derive the following equation:

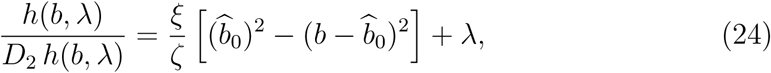

where *D*_2_ *h >* 0 (Lemma 1). Hence, the roots 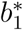 and 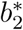of *h*(·, *λ*) are given by

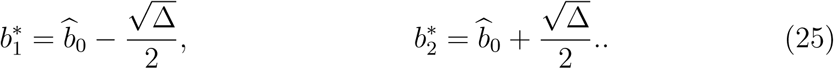

Observe that 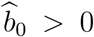 if and only if *D*_1_ *h*(0, 0) *>* 0 according to (14). From the definitions of Δ and equation (15), the following equations are equivalent:

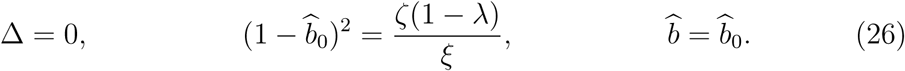

If one (hence all) of the equations in (26) is true, then 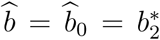 by (25) and equivalently 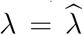 by Lemma 3. We may write 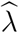 in terms of the dimensionless parameters by solving for Δ = 0:

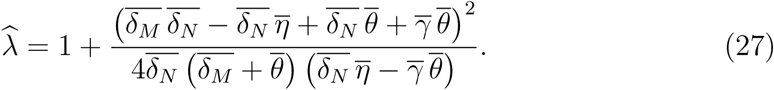

### Theorem 3.

*Suppose that 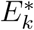 exists for a given k, and that all eigenvalues of 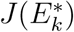 have nonzero real part. Then 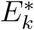 is a saddle point for k* = 1 *and asymptotically stable for k* = 2.

*Proof.* Consider the Jacobian matrix 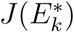. To obtain a practical expression of 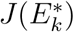, we simplify the first and second diagonal entries by application of equations (20) and (21), respectively; by *i*th diagonal entry, we mean the (*i, i*)-entry. For the off-diagonal entries, we perform the following algebraic manipulations:

(i) In the top row, write 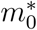 in terms of 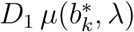.

(ii) In the middle row, evaluate 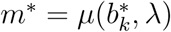 and write 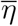 in terms of 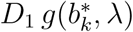.

(iii) In the bottom row, write 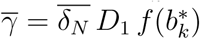.

Additionally, we apply the equation 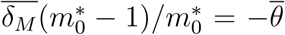 wherever simplification is desired. Thus, we arrive at the following expression:

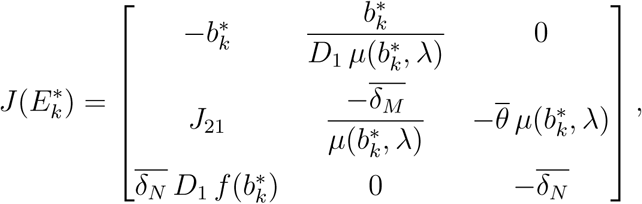

Where

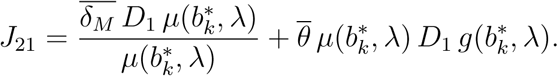

Denoting the trace and determinant by tr and det, respectively, 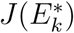 must satisfy three Routh-Hurwitz conditions for 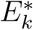 to be asymptotically stable. The first condition holds for both 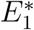 and 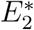, that is:

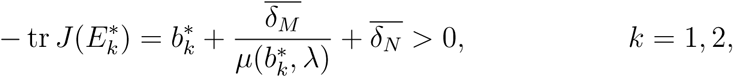

given that *µ >* 0. The second condition requires

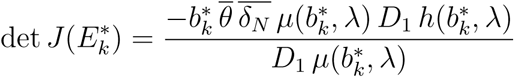

to be negative. Since *D*_1_ *µ <* 0, the determinant det 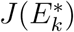 shares the same sign with 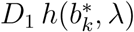. Thus, by Lemma 3, the second condition fails for 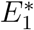 because 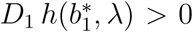. Since we assumed that all eigenvalues have nonzero real part, 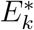 must be a saddle point for *k* = 1. Meanwhile, Theorem 2 implies that for 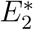 to exist, it is necessary that either *λ >* 0 or 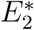 coexists with 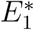. In either case, 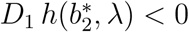 and the second Routh-Hurwitz condition holds for *k* = 2.

We are left to verify the following last Routh-Hurwitz condition for 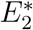, *i.e.*, assuming that *k* = 2:

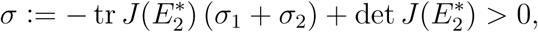

Where

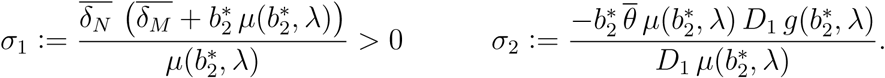

Assuming that 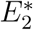 exists, observe that *σ*_2_ has the same sign with 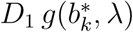. Since 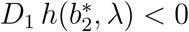 and *f* has a positive first derivative (Lemma 1), we have

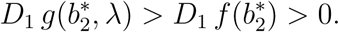

and *σ*_2_ *>* 0. Moreover, we have

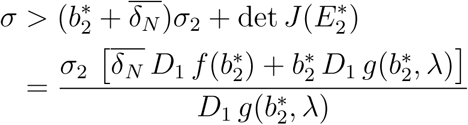

and *σ >* 0. Therefore, 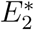 satisfies all three Routh-Hurwitz conditions and is consequently asymptotically stable.

The existence of roots, as well as their stability—corollary of Theorem 2, and Theorem 3, respectively—are summarized in Figure 4, showing an illustration of the different stability regions in the 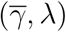-plane, and numerical bifurcation plots for *b* as a function of *λ*.

**Figure 4:**
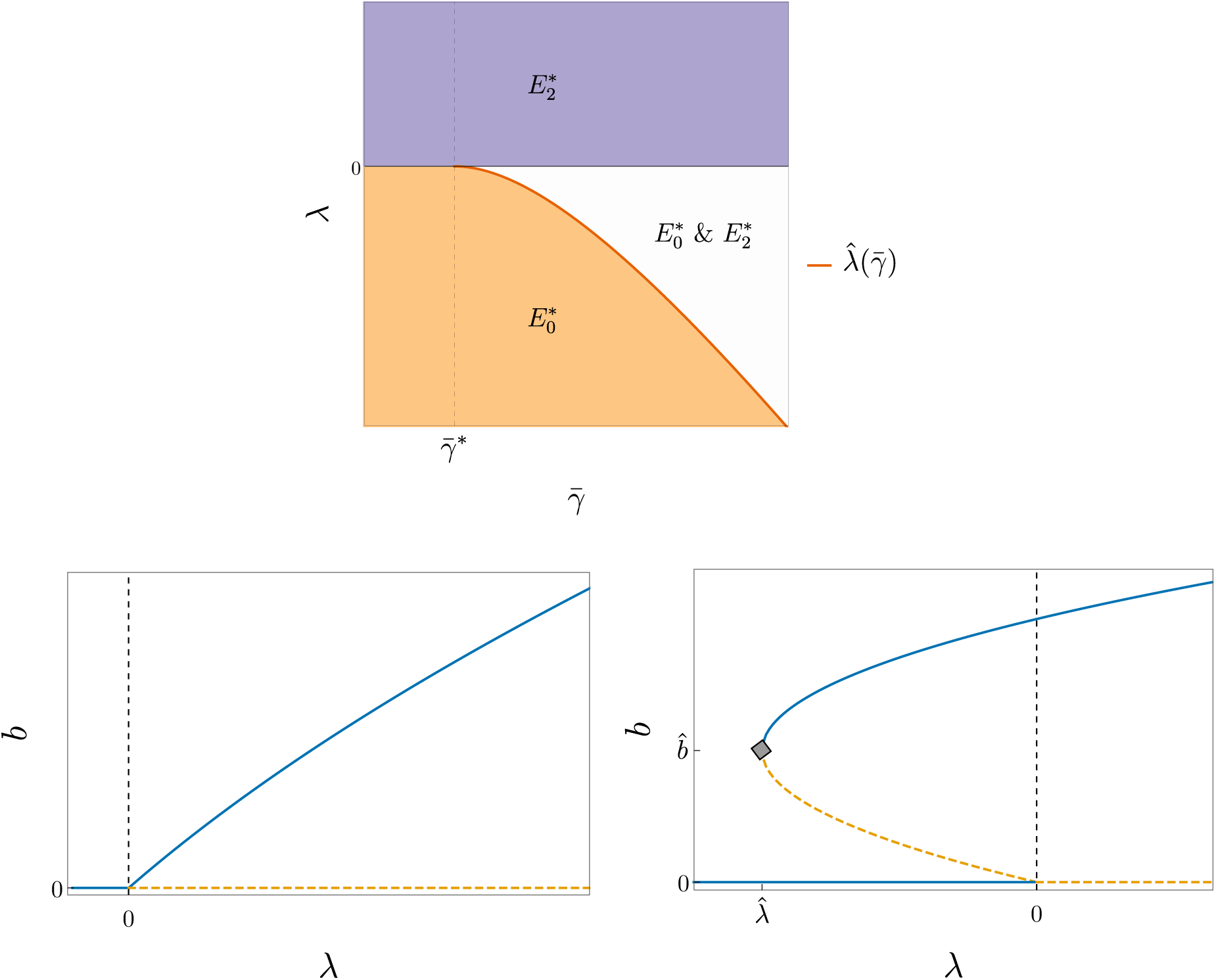
**Top:** Regions of stability determined by *λ* and 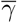, each highlighting the corresponding stable equilibrium points; 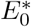 has *b* = 0, while 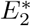 takes the largest steady-state value 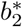 for *b*. The specific value 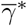 of *γ* is given in equation (19), while 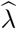 is explicitly given in (27).

**Bottom:** Bifurcation diagrams of the system (4)—(6) without (left) and with (right) bistability— respectively, 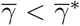 and 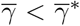. To generate the diagrams, all parameters were fixed except for 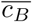, which was obtained for a given *λ* from (7). Blue, solid lines: stable equilibria; orange, dashed lines: unstable equilibria. The gray diamond appearing in the right panel highlights the bifurcation point 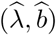.

## Discussion

Our study is centered on the problem of identifying biological factors that contribute to the elimination of the pneumococcus (*Streptococcus pneumoniae*). Experiments by Smith et al. [19] suggested that inoculum size (dosage) determines the outcome of bacterial clearance or persistence. In this case, groups of mice were infected with the pneumococcus at different dose sizes. Each group corresponded to a single bacterial outcome indicated by the titer readings, depending on whether the dose size is above or below a threshold. In contrast, experiments from Duvigneau et al. [27] showed that inoculum size is not the only factor contributing to bacterial clearance. Here, all mouse subjects were given doses of identical size. Towards the end of the experiment, the bacterial load was undetectable in some mice and sustained in the rest (Figure 2). The murine experiments of Smith et al. [19] and Duvigneau et al. [27] provide distinct perspectives on the trade-off between dosage and bacterial fate, restricted by the microbial instances compatible with the experimental design.

By modeling bacterial kinetics with generalized innate-regulatory immune responses, our mathematical analysis reveals a qualitative trichotomy of this trade-off that acknowledges the experiments of both Smith et al. [19] and Duvigneau et al. [27]. Indeed, our model, given by system (1)–(3) and the equivalent dimensionless form (4)–(6), may be considered an abstraction of previous formulations [17, 18, 16, 15, 19] where overall immune responses are considered instead of specific phagocyte populations and chemical mediators. The trichotomy is given by the following cases: (i) *dose-independent clearance*, where the immune response clears the pneumococcus independent of dose size; (ii) *dose-independent persistence*, where the pneumococcus outgrows immunity regardless of initial dose; and (iii) *dose-limited clearance*, where the immune system successfully eliminates the pneumococcus only in small quantities. Cases (i) and (ii) are corroborated by Duvigneau et al. [27], whereas case (iii) is supported by Smith et al. [19].

We remark that successful clearance of the pneumococcus may also be attributed to empirical characteristics of the infection other than the inoculum size. Mochan et al. [21] formulated a model validated with murine datasets including [19] to describe pulmonary and extrapulmonary pneumococcal kinetics with total phagocyte levels and damage to epithelial cells (cellular debris) and a homogeneous population of activated phagocytes. The corresponding simulations indicate that phagocyte clearance efficiency varies between mouse strains. A follow-up study [22] using the same model shows that mutations in the pneumococcal strain can influence transient reduction in pneumococcus. In addition, Schirm et al. [20] adapted the monocytederived macrophage murine model in [19] and incorporated inhalation and antibiotic effects to pneumococcal growth and elimination. The models of Mochan et al. [21, 22] and Schirm et al. [20] are directed towards investigating the effects of treatment and strain variations on pneumococcal clearance. However, their modeling approaches focus on validation with experimental data instead of bifurcation. While our mathematical model only considers bacterial kinetics with generalized innate and regulatory levels via three equations, the results of our comprehensive stability analysis could provide valuable and testable hypotheses regarding pathogen clearance.

The following quantities were determined to drive the dynamics of our model:

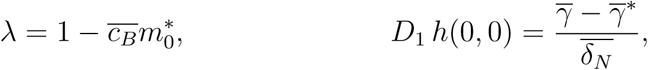

where

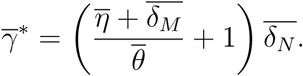

See equations (7) and (19). The stability of the bacteria-free steady state 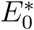 according to *λ* (Theorem 1) portrays the effectiveness of the innate immune response to clear small pneumococcal quantities. Now, *λ* is a linearly decreasing function of 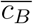, which is proportional to the clearance rate *c*_*B*_ with the initial innate response level *M*_0_ = *M*(0) and all other parameters fixed. Since 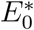 is stable when *λ <* 0 or 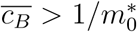 (Theorem 1), a rapid innate clearance (large *c*_*B*_) could promote bacterial eradication at small quantities with the goal of making 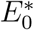 stable. Effective clearance may also hold in mild conditions (moderate values of *c*_*B*_ and *M*_0_) for a large innate level 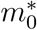. By the same token, 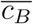 exhibits inverse proportionality with the maximum logistic proliferation rate *r* of the bacteria. Thus, the outgrowth of the pneumococcus may benefit from rapid proliferation (large *r*).

The overall dynamics of our model follows from our main results for positive equilibria. The corollary of Theorem 2 determines which of 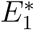 and 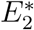 exist, and Theorem 3 establishes that 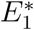 is unstable and 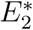 is asymptotically stable. Since *D*_1_ *h*(0, 0) has the same sign with 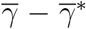, we may frame our discussion in terms of the dimensionless parameter 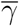. As illustrated in Figure 4, 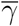 determines which of the three bacterial outcomes (clearance, persistence, dose-limited clearance) are possible while *λ* decides which outcome the model predicts.

If the model assumes that 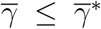, then bistability does not occur and the model only predicts dose-independent clearance and persistence. That is, the stable equilibrium point is uniquely given by 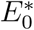 for *λ <* 0 and 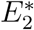 for *λ >* 0. Hence, we expect the innate immune system to eliminate the bacteria for *λ <* 0, and for the bacteria to persist for *λ >* 0, regardless of the initial bacterial concentration. As 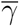 increases so that 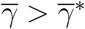, the negative values for *λ* corresponding to dose-independent clearance are restricted to the interval 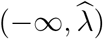 where 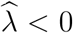. Moreover, bistability holds for all values of *λ* in the interval 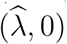, where the unstable equilibrium point 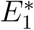 coexists with the stable equilibria 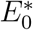 and 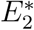. The size of the interval 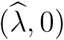 changes with 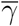 according equation (27). We emphasize that the monostability at 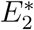 when *λ >* 0 is independent of 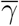.

When scaled with 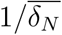 (with fixed 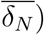, we find that 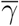 is directly proportional to the tissue carrying capacity *K*_*B*_ and the ratio *γ/*(*δ*_*N*_ *N*_0_) of innate response promotion to constant replenishment rate. Hence, the model predicts that abundant tissue resources (hypothetically large *K*_*B*_), and rapid activation of regulatory responses (*γ > δ*_*N*_ *N*_0_) contribute to the range of parameter values for bistability and dose-limited clearance. In light of the monostability of 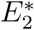 above, we predict that the corresponding dose-independent persistence may neither depend on tissue carrying capacity nor the activation of regulatory responses.

In the bistability case, the bacterial load 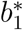 at 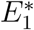 can serve as a threshold for bacterial clearance. Based on our bifurcation diagrams (Figure 4), one could naively deduce that the immune system clears the bacteria for 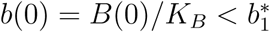, and the bacteria succeeds in colonizing the host when 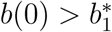 (cf. [25]). However, we must emphasize that our model is a *three*-dimensional system where stable manifolds of a saddle node may be one-dimensional curves or two-dimensional surfaces. A deeper analysis requires investigating the stable and unstable manifolds of 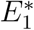, which delimit the basins of attraction for 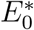 and 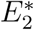.

At this point, we discuss generalizations and future directions of our work. The aforementioned local stability as dependent on *λ* may be qualitatively identified with compatible systems exhibiting nonlinear interaction terms. This can be achieved with the function

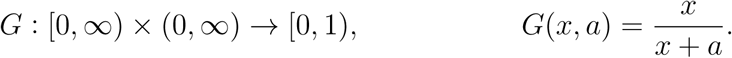

For a fixed *a*, the function *G*(*·, a*) typically introduces saturation effects on a growth/decay rate: the value of *G*(*x, a*) approaches its upper bound (*G≈*1) with larger values of *x*. In different biological contexts, *G* is associated with the Monod growth term for microorganisms, the Michaelis-Menten equation for enzyme kinetics, and the Holling Type-II functional response for predator-prey dynamics.

To demonstrate the robustness of our results to nonlinear interaction terms, we show in Figure 5 that a modification of the model (1)–(3) incorporating nonlinear interaction terms yields comparable qualitative dynamics. Moreover, the same trichotomy of bacterial outcomes applies here. The modified system is given by

**Figure 5:**
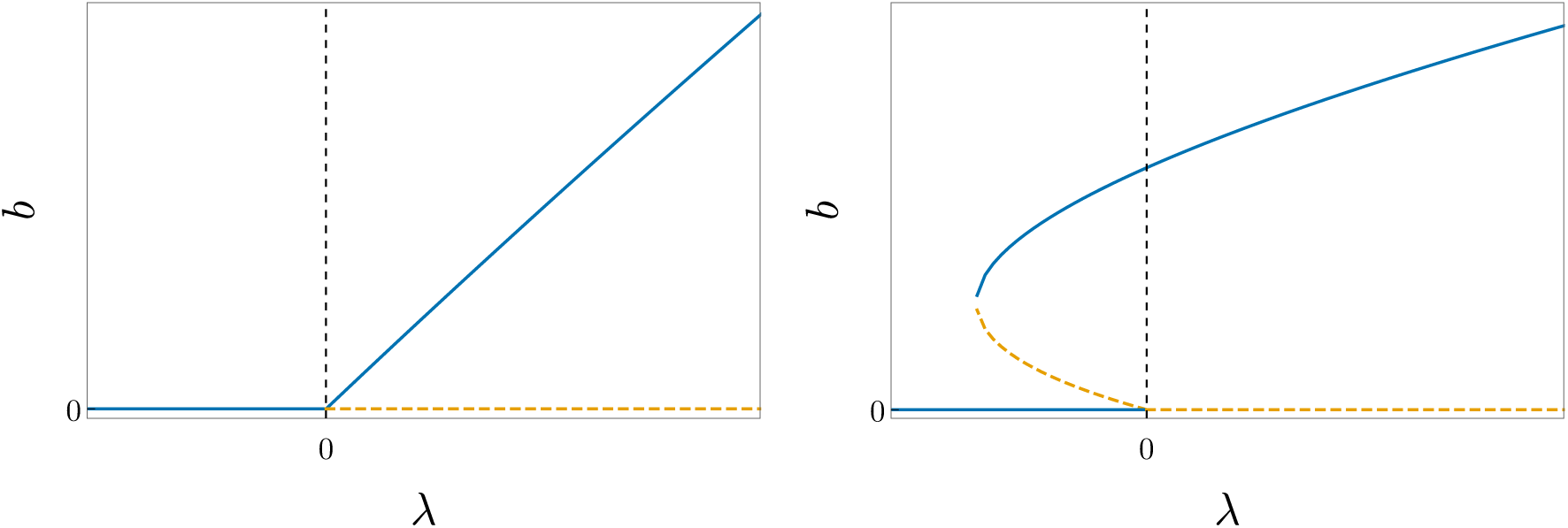
Bifurcation diagrams generated by the system (28)–(30), where *b* = *B/K*_*B*_ and *λ*, given by (31), is the eigenvalue that determines the stability of the unique equilibrium point satisfying *b* = 0. These diagrams are generated via the same procedure as the ones from Fig. 4, and are shown for 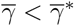, corresponding to the case without bistability (left), and 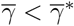, corresponding to the bistable case (right); here, 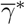 is given by (32). Blue, solid lines: stable equilibria; orange, dashed lines: unstable equilibria.

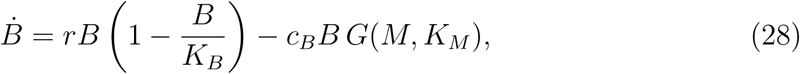

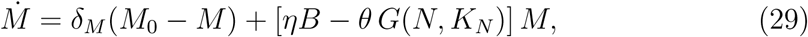

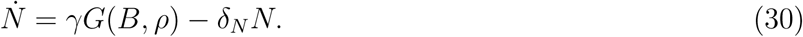

This model imposes the following effects: (i) saturated bacterial clearance with high levels of innate immune response; (ii) bounded regulation of innate response; and (iii) limited increase in regulatory levels for larger bacterial loads.

Proceeding as before, we generate a bifurcation diagram for (28)–(30) based on the eigenvalue

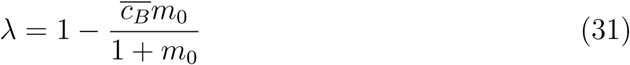

characterizing the stability of the unique bacteria-free equilibrium. Here, we have used the dimensionless quantities *m*_0_ = M_0_/*K*_*M*_ 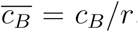, and the time has again been rescaled as *τ* = *t/r*. The bistability region is now determined by

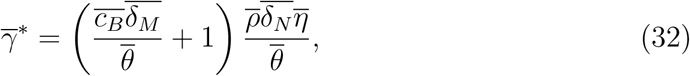

with 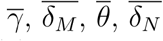 and 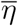 as defined before, and 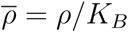. As in the original model (1)–(3), the parameter 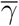 determines whether bistability exists, and *λ* determines which outcome is predicted. However, the formulation of *λ* in (31) is independent of parameters pertaining to innate response as opposed to the original formulation in (7). This observation suggests that, to an extent, bacterial clearance may depend on other factors aside from innate response when nonlinear cellular responses are in action.

## Acknowledgements

This work was supported by the Boehringer Ingelheim Stiftung (Exploration Grant, VIBA project) and the Alfons und Gertrud Kassel-Stiftung.

